# Mutualistic interactions between *B. subtilis* and seeds dictate plant development

**DOI:** 10.1101/2021.06.07.447311

**Authors:** M.V. Berlanga-Clavero, C. Molina-Santiago, A.M. Caraballo-Rodríguez, D. Petras, L. Díaz-Martínez, A. Pérez-García, A. de Vicente, P. C. Dorrestein, D. Romero

## Abstract

A tightly coordinated developmental program controls precise genetic and metabolic reprogramming that dictates efficient transition of the seeds from dormancy to metabolically active seedlings. Beneficial microbes are known to stimulate the germination of the seeds or adaptation of the seedlings; however, investigations of exact mechanisms mediating these interactions and the resulting physiological responses of the plants are only beginning. *Bacillus subtilis* is commonly detected in the plant holobiont and belongs to the group of microbes that provide multifaceted contribution to the health of the plants. The present study demonstrated that *B. subtilis* triggered genetic and physiological responses in the seeds that determined subsequent metabolic and developmental status of adult plants. Chemically diverse extracellular matrix of *Bacillus* was demonstrated to structurally cooperate in bacterial colonization of the seed storage tissues. Additionally, an amyloid protein and fengycin, which are two components of the extracellular matrix, targeted the oil bodies of the seed endosperm, provoking changes in lipid metabolism or accumulation of glutathione-related molecules that stimulated two different plant growth programs: the development of seed radicles or overgrowth and immunization of adult plants. We propose this mutualistic interaction is conserved in Bacilli and plant seeds containing storage oil bodies.

## Introduction

In a scenario of constantly increasing demand for sustainable agriculture, the success of seed germination and seedling development of crops determine boosted productivity and avoid economy- and resource-related problems^1,2^. Seed germination is a highly complex biological process tightly controlled by interconnected hormone-related pathways that ensure efficient mobilization of the nutrients leading to radicle emergence and subsequent proliferation of adult plants^3^. Improvement of these early stages of plant development is an important challenge for the success of environmentally friendly strategies based on the use of beneficial microbes. Seed inoculation with plant growth-promoting rhizobacteria (PGPR) is a common procedure to ensure bacterial colonization of the plants and beneficial effects of PGPR after germination^4^. Better understanding of the exact mechanisms that direct these specific host-microbe interactions and consequences at the growth-promoting level are important to ensure feasible and robust treatments.

*Bacillus subtilis* and closely related species mutualistically coexist with the plants, providing a plethora of beneficial actions, such as biofertilization or protection against biotic and abiotic stresses^5^. Ecological traits contributing to bacterial fitness and bioactivity in the plants are due to polyvalent secondary metabolites, sporulation, or biofilm formation^6^. Biofilms are formed by the cells embedded in a self-secreted extracellular matrix (ECM), which is a chemically complex megastructure consisting of proteins, exopolysaccharides, nucleic acids, and secondary metabolites^7–9^. The ECM is known to be essential for efficient colonization of the plant organs^10,11^, and some secondary metabolites mediate bacterial communication with the plants by triggering physiological responses associated with defense or growth^12,13^. The present study investigated the mechanism by which seeds respond to stimulatory activity of *Bacillus* and exact functionality of the ECM in this mutualistic microbe-host interaction. The results demonstrated that the ECM components cooperatively contributed to colonization of the seeds; however, only lipopeptide fengycin and amyloid protein TasA mediated chemical dialogue resulting in two different plant responses: i) promotion of radicle growth after seed germination or ii) overgrowth of adult plants and protection against the aerial necrotrophic fungus *Botrytis cinerea*. Both molecules targeted the lipid storage oil bodies of the seed endosperm, leading to specific metabolic reprogramming early after seed germination responsible for one of these two responses. Thus, the mechanism by which seeds respond to the presence of *Bacillus* cells was influenced by the nutrient storage of the seeds and internal anatomy.

## Results and discussion

### Bacterization of the seeds with *B. subtilis* dictates metabolic reprogramming and overgrowth of adult plants

Beneficial rhizobacteria can promote seed germination that involves two different and genetically controlled stages to ensure further growth of adult plants in a supportive environment: germination *sensu stricto* and growth of the emergent radicle^1^. The phenotype derived from the seeds bacterized with *B. subtilis* resulted in larger radicles compared with those grown from untreated seeds (Fig. 1A). In contrast, the germination rates (initial emergence of the radicle) of melon seeds bacterized with the cells of *B. subtilis* NCIB3610 did not change compared with that of untreated seeds; consistently, the expression levels of *GA20ox1* and *CYP07A1*, which are the two genes involved in the modulation of the determinant ratio of abscisic acid and gibberellins^14^, were statistically similar in the treatment groups (Extended Data Fig. 1a, b). Thus, complete transcriptomic analysis of the seeds 16 hours after the treatment with the bacterial inoculum did not detect any changes in the expression levels of the genes involved in the germination-related hormone signaling pathway. The presence of 141 differentially expressed genes (DEGs) suggested a very specific response of the plants to bacterial treatment. Accelerated activation of seed metabolism was sustained by the induction of the genes involved in carbon metabolism and photosynthesis and strong repression of heat shock proteins and structural proteins of lipid storage vesicles (oleosin and caleosin), which may reflect a faster decline in these transcripts during germination^15–17^ (Fig. 1B, Extended Data Fig. 1c, d). This optimized metabolic activity reasonably explained an increase in the total area of the radicles produced by bacterized melon seeds (Fig. 1A). Notably, this post-germinative stimulatory activity was long–lasting considering that adult plants emerged from bacterized seeds developed a higher number of the roots and more vigorous canopies than those developed by the untreated plants (Fig. 1C).

**Figure 1.**
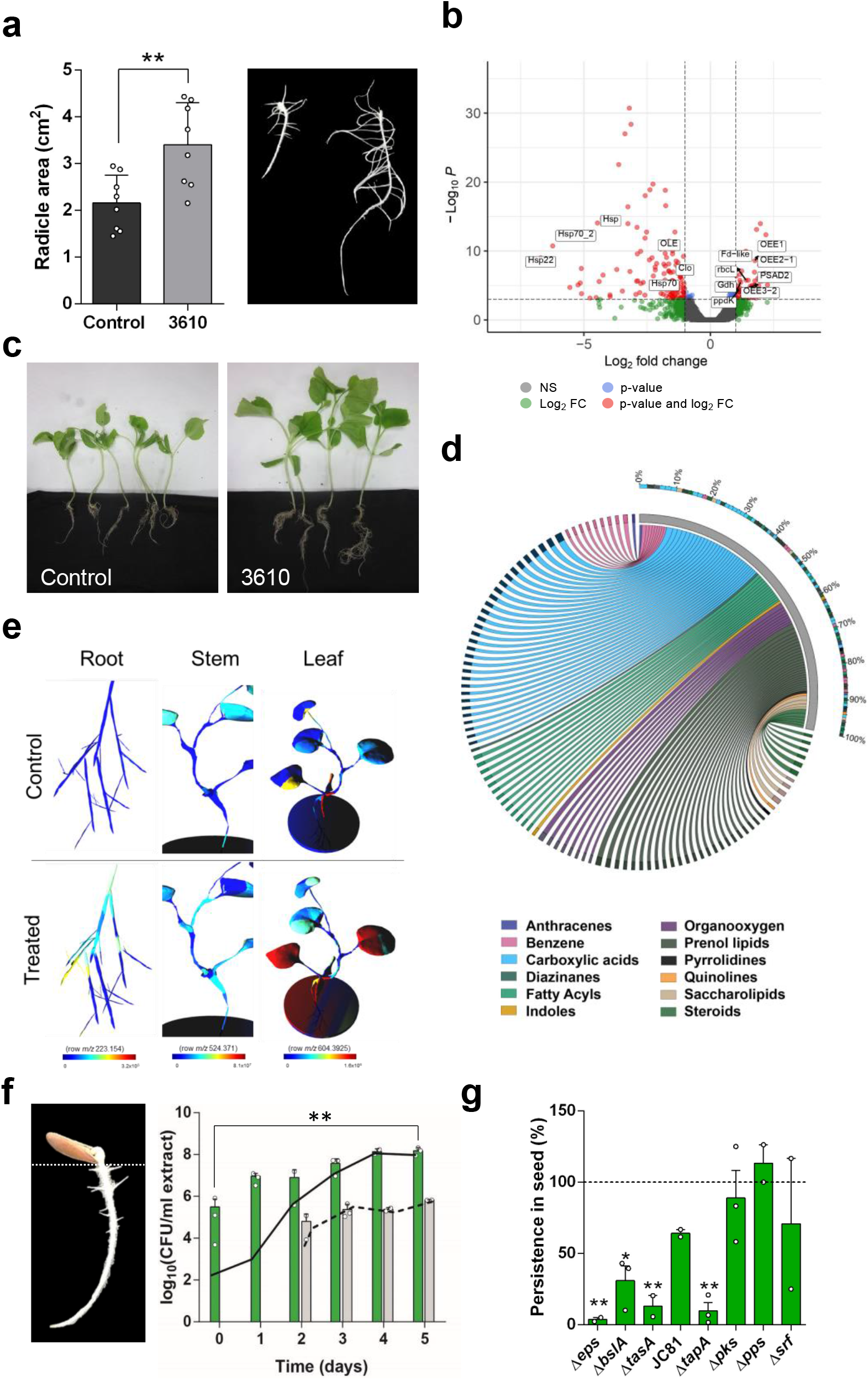
Interaction of *B. subtilis* with the seeds stimulates radicle development and results in overgrowth of adult plants. **A** Left: Average radicle areas after seed treatments with *B. subtilis*. Error bars represent SD. Statistical significance was assessed by a t test (n= at least 7). Right: representative radicles from a *B. subtilis*-treated seed (right) and untreated seed (left) five days after the treatments. **B** Volcano plot of DEGs identified by RNA-seq in bacterized seeds and untreated seeds 16 hours after the treatment. Tags label the genes related to seed germination progress: OEE1 (oxygen-evolving enhancer protein 1), OEE3-2 (oxygen-evolving enhancer protein 3-2, chloroplastic), OEE2-1 (oxygen-evolving enhancer 2-1, chloroplastic), Fd-like (ferredoxin-like), PSAD2 (photosystem I reaction center subunit II, chloroplastic), Gdh (glutamate dehydrogenase), rbcL (ribulose bisphosphate carboxylase small chain), ppdK (pyruvate, phosphate dikinase), pckG (phosphoenolpyruvate carboxykinase), Clo (caleosin); Hsp70 (heat shock 70 kDa protein), Hsp (class I heat shock protein), Hsp70 (heat shock 70 kDa protein_1), and Hsp22 (22.0 kDa class IV heat shock protein). **C** Adult plants grown from the seeds treated with *B. subtilis* (3610, right) or from untreated seeds (control, left). **D** Circos plot of 100 metabolites more abundant in the leaves of the plants grown from bacterized seeds than in the leaves of the control plants. **E** Distribution of three representative features classified as lipids with significant differential abundance between the regions of the plants emerged from untreated seeds (control) and *B. subtilis-treated* seeds. **F** *B. subtilis* dynamics (CFU counts) in the seed and radicle extracts during the first five days after seed treatment. Bars represent average values with error bars (SEM) of total CFU in the seeds (green bars) and radicles (gray bars), and continuous and discontinued lines represent CFU corresponding to the number of spores in the seeds and radicles, respectively. Statistical significance was assessed by two-tailed independent t-tests between initial and final time points (n= at least 2). **G** Bacterial persistence of the ECM mutants relative to that of the WT mutant (assumed to be 100%) in the seed extracts five days after seed treatment. Average values and error (SEM) are shown. Statistical significance was assessed by after one-way ANOVA with I Dunnett’s multiple comparisons test (each treatment vs. WT treatment) (n= at least 2).

Interactions of PGPR with adult plants lead to the accumulation of certain plant metabolites associated with the promotion of plant growth, adaptation to abiotic stress, or defense against microbial diseases^18^. Thus, we analyzed the metabolome of adult plants that emerged from bacterized seeds to define putative major metabolic changes correlated with the long-lasting growth-promoting effect. Adult plants derived from treated or untreated seeds were sectioned into the roots, stems, and leaves and analyzed by LC/MS/MS and Feature Based Molecular Networking analyses (FBMN) using the GNPS platform^19^. Statistical analyses of the data obtained using all regions indicated that carboxylic acids, lipids, and lipid-like molecules were predominant chemical classes of metabolites differentially detected in aerial regions of the plants (Fig. 1D and Extended Data Fig. 2), and metabolomic composition of the roots was not significantly different between the groups. Changes in fatty acyls, glycerophospholipids, steroids, and prenol lipids were considered the major metabolic signature referring to lipids of the plants grown after bacterial treatment of the seeds (Fig. 1E, Extended Data Fig. 3a, b). Induction of certain secondary metabolites produced by the plants as a result of their interaction with PGPR have attracted the most interest because of versatile functionalities and biotechnological applications of these compounds^20,21^. However, other molecules, such as the organic acid malate, key amino acids related to nitrogen assimilation, fatty acids, and hydroxycinnamic acid derivatives, have been detected in various cases of plant-PGPR interactions and considered biomarkers of PGPR priming^21,22^. Tryptophan-derived secondary metabolites have been recently shown to be related to the key genes involved in general nonself response to bacterial pathogens^23^. The results of metabolomic analysis of adult plants revealed differential accumulation of L-tryptophan and cinnamic acid, both belonging to the top 50 metabolites more abundant in the leaves of the plants grown from the treated seeds (Extended Data Fig. 3c).

### Complementary contribution of the ECM to *Bacillus* ecology and the promotion of seed radicle growth

Our previous findings suggested that bacterization of the seeds with *B. subtilis* cells potentiates metabolomic changes with relevant implications for the development of adult plants. Initially, we investigated the dynamics of the bacterial population of *B. subtilis* five days after seed bacterization in two different regions, including the seed and emergent radicle (Fig 1F). In the radicles, the size of the bacterial population (gray bars) remained unchanged, and the bacteria remained entirely sporulated (discontinuous lines) from the very first moment of radicle emergence (2 days after inoculation). However, the bacterial population was significantly increased in the seeds (green bars) over five days of the experiment and became almost entirely sporulated (continuous lines) four days after bacterization. These results suggested that *B. subtilis* cells, mostly spores, were passively dragged by the emergent radicle; instead, these cells could have colonized and proliferated inside the seeds. The results of scanning electron microscopy analysis (SEM) of bacterized seeds confirmed bacterial colonization of the inner regions of the seeds. The results of confocal laser scanning microscopy (CLSM) analysis of transversely sectioned seeds previously bacterized with fluorescently labeled *B. subtilis* (CellTracker™ cm-DII) indicated the accumulation of bacterial cells in the storage tissues near the seed micropyle, which is the natural entry point of bacteria into the seeds^24^ (Extended Data Fig. 4a, c). The cell density of the WT strain and *Δhag* (flagellum), Δ*eps*, or Δ*srf* mutant strains, which are known to have altered swimming, sliding, or swarming motility, respectively^25–27^, was essentially similar in the whole seeds or in two differentiated parts inside the seeds, including the micropylar and opposite chalazal regions, four hours after the treatment (Extended Data Fig. 4b). Overall, these findings indicated that the growth-promoting activity may be triggered by bacterial cells that entered and colonized the seed storage tissues, which is a process that did not appear to rely on a specific type of bacterial motility.

Biofilms are a known stage of bacterial life cycle; the cells in a biofilm are embedded in a chemically complex and functionally versatile extracellular matrix (ECM) whose components complementarily participate to benefit the community^7,28,29^. A number of studies described multifaceted contributions of biofilms to ecology of bacterial cells in their interactions with the hosts (adhesion, colonization, persistence, or protection)^11,30–32^ or to protection against pathogens either in the rhizosphere or in the phyllosphere^10,33,34^. Thus, we hypothesized that the ECM provides a relevant contribution to the beneficial *Bacillus*-seed interactions. As suggested by the structural functions, the cell density of single mutants ΔtasA, ΔtapA, Δeps, or ΔbslA was significantly decreased compared with that of the wild type bacteria (assumed to be 100%) five days after seed bacterization (Fig. 1G). However, this result was not associated with the attachment of bacterial cells to the seeds a few hours after the treatments (Extended Data Fig. 5a). A similar bacterial population pattern was monitored in the emergent radicles, which, as described above in the case of WT, may reflect dragging of bacterial cells, which have initially entered the seeds, by the growing radicles (Extended Data Fig. 5b). The Δsrf, Δpks, or Δpps strains manifested arrested production of secondary metabolites surfactin, bacillaene, or fengycin, respectively (also belonging to the ECM); however, these strains did not deviate from the population dynamics observed in the case of the WT strain (Fig. 1G and Extended Data Fig. 5). A decrease in the bacterial cell population suggested that the majority of ECM mutations led to a significant decrease in the percentage of an increase in radicle growth compared to that detected after the treatment with the wild-type strain (Fig. 2A). However, the Δsrf strain retained radicle growth-promoting activity similar to that of the WT strain; this finding along with similarities in the levels of persistence with those of the WT strain indicated that surfactin was not implicated in the growth-promoting activity. In contrast to a correlation between bacterial cell density with plant growth promotion, the poorly persisting ΔtasA strain retained the promoting activity, and the Δpks and Δpps strains failed to promote radicle growth despite considerable persistence on the seeds (Fig. 2A).

**Figure 2.**
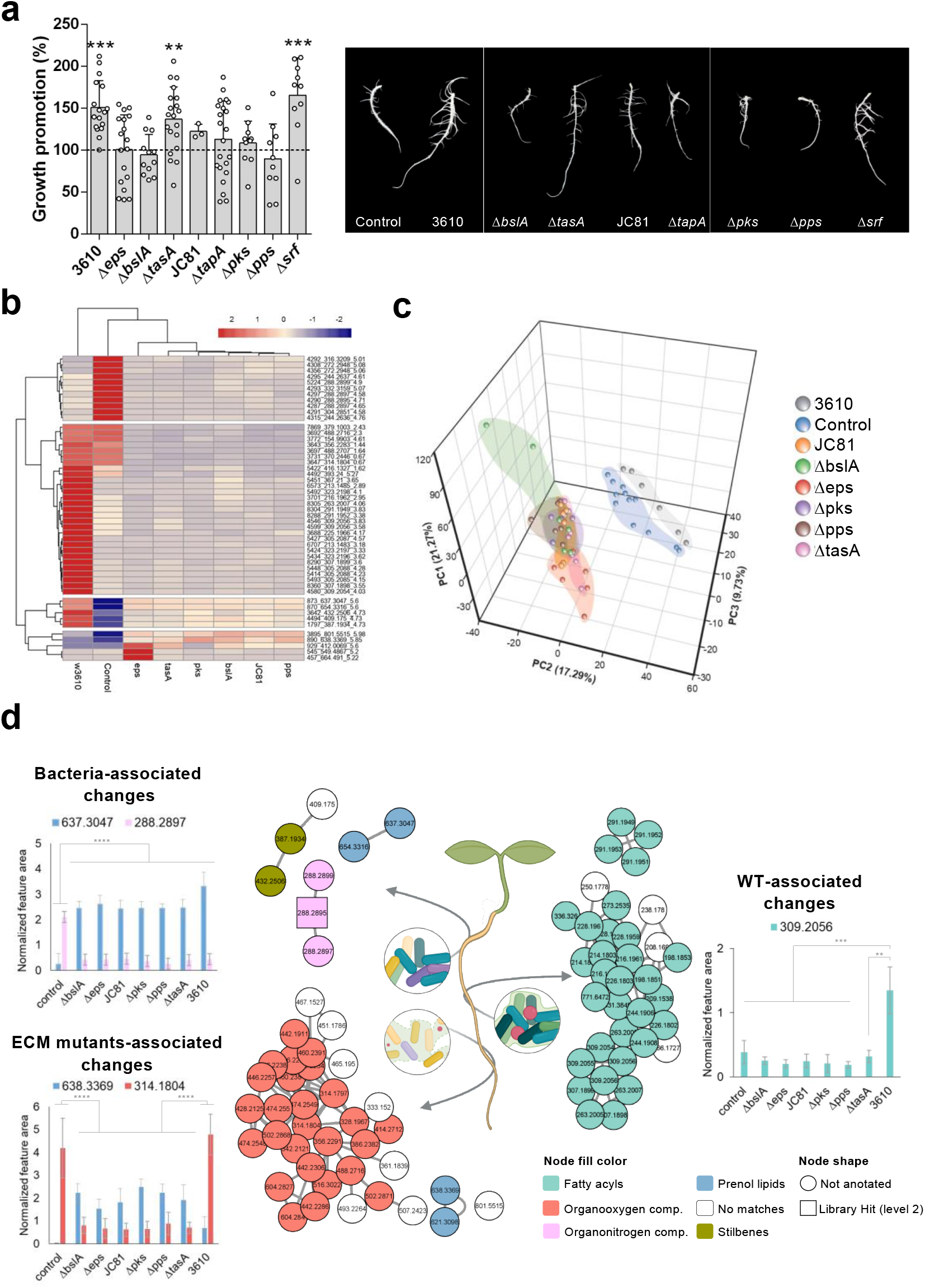
Components of the extracellular matrix of *B. subtilis* trigger metabolic reprogramming of the seed radicles related to plant growth stimulation. **A** Left: percentage of radicle growth increase in the seeds treated with wild-type or ECM mutant cells (five days after seed treatment) normalized to the average radicle area of untreated seeds (100%, discontinued line). Average values are shown, and error bars represent SD. Statistical significance was assessed by one-way ANOVA with *post hoc* Dunnett’s multiple comparisons test (each treatment vs. control treatment) (n= at least 10 except JC81 with n=3). Right: Representative radicles from untreated seeds and seeds treated with wild-type or ECM mutant cells five days after the treatment. **B** Heatmap of the hierarchical clustering of the top 50 features of impacted molecular families in the radicles from bacterized seeds. **C** PCA 3D score plot of the metabolome of radicles clusters the samples based on the presence, absence, or alteration of the ECM after bacterial treatments. The percentage of variation explained by each principal component is indicated on the axes. **D** Network analysis of representative features related to the presence of bacteria or ECM. Normalized abundances in the radicles in seeds subjected to various treatments are represented in features of all groups. Average values are shown with error bars representing SD. Statistical significance was assessed by one-way ANOVA with *post hoc* Dunnett’s multiple comparisons test (n=3).

Recently, the absence of the amyloid protein TasA, in addition to influencing biofilm formation, has been reported to provoke important alterations in cellular physiology, including overproduction of fengycins^11^, which appears to be implicated in the promotion of radicle growth (Fig. 2A). Thus, to clarify the contribution of TasA as a structural ECM component to the growth-promoting activity, we tested the JC81 strain, which expresses a version of TasA that fails to fully restore the biofilm formation phenotype and reverts the physiological status of the cells to the levels comparable to that of the WT strain^11^. JC81 failed to promote seed radicle growth but persisted on the seeds at a level comparable to that of the WT, Δpks, or Δpps strains (Fig. 1G and 2A). These results demonstrated two distinctive and complementary contributions of the ECM components to the promotion of seed radicle growth: i) EPS, BslA, and TapA ensured the persistence of bacterial cells in the seeds; ii) bacillaene (*pks*) and fengycin (*pps*) mediated the chemical dialogue of *Bacillus* with the seeds; and iii) amyloid TasA was involved in both activities.

### Different integrity states of the bacterial ECM induce distinct metabolic changes in emerged radicles

To establish a connection between the chemical dialogue of *Bacillus* with the seeds and the overgrowth phenotype of adult plants, we characterized the metabolic signatures defining the stimulatory effects on radicle growth at the initial stages after seed germination. We analyzed and compared the metabolic statuses of radicles five days after the treatment with the cells of the WT or various ECM mutant strains by LC/MS/MS and FBMN analyses^35^. Hierarchical clustering using Metaboanalyst was used to visualize the top 50 features impacted by the treatment that distinguish the samples treated with bacteria from untreated samples or WT, Δeps, or ECM mutant strain-treated samples from other samples (Fig. 2B). The results of principal component analysis (PCA) emphasized a similarity of the distributions of the treatments with ECM mutant strains, and the samples treated with the WT strain and untreated samples formed independent groups (Fig. 2C). Previous classification was combined with FBMN analysis and *in silico* chemical class analyses (Classyfire^36^ and MolNetEnhancer^37^) identified specific molecular families belonging to these groups according to the abundance patterns and associations of the patterns with specific chemical classes (Fig. 2D). The changes associated with the treatment with bacteria included a decrease in organooxygen compound analogs of sphinganines and an increase in reduced abundance of the prenol lipid and stilbene molecular families. The presence of a functional ECM in the WT strain triggered the accumulation of fatty acyls belonging to two different clusters, and treatments with ECM mutants induced a clear decrease in organooxygen compounds and accumulation of specific prenol lipids. These results demonstrated a specific metabolic response of the plants to chemical features of the ECM of *B. subtilis*.

### *Lyso*phosholipids and glutathione are key metabolites involved in different growth-promoting effects of TasA and fengycin

All previous experiments suggested that TasA and fengycin are the most relevant molecules of the ECM mediating the interkingdom chemical dialogue of *Bacillus* cells with the seeds. TasA is a polymorphic protein able to adopt a variety of structural conformations, from monomers to aggregates or insoluble fibers, in response to physicochemical variations of the environment. These morphotypes are biochemically different, and studies on this protein and other amyloids have revealed that they perform distinct biological functions^11,38,39^. Polymerization of purified homogenous TasA monomers was triggered and its ability to promote seed radicle growth was tested at various stages of aggregation (Extended Data Fig. 6a). The results confirmed that purified TasA was able to promote seed radicle growth, and this effect relied on large aggregates of the protein. Moreover, fresh apoplast fluid extracted from the melon seeds promoted the polymerization of TasA to form aggregates (Extended Data Fig. 6b), indicating that this largely active polymerized form of TasA was predominant inside the seeds. Evaluation of the stimulatory activity showed that 3 μM solution of the most active form of TasA or 10 μM solution of purified fengycin significantly increased the area of the radicles compared with that in untreated seeds (Fig. 3A). The lack of stimulatory activity of purified BslA or surfactin confirmed our previous finding obtained using single mutants arrested in the production of all these molecules (Fig. 2A) and demonstrated that additions of exogenous protein or secondary metabolites did not necessarily manifest growth-promoting effects (Fig. 3A). Iturin is not produced by *B. subtilis*; however, purified commercially available iturin failed to promote radicle growth after seed treatment, indicating that lipopeptides do not have a universal promoting effect (Extended Data Fig. 7).

**Figure 3.**
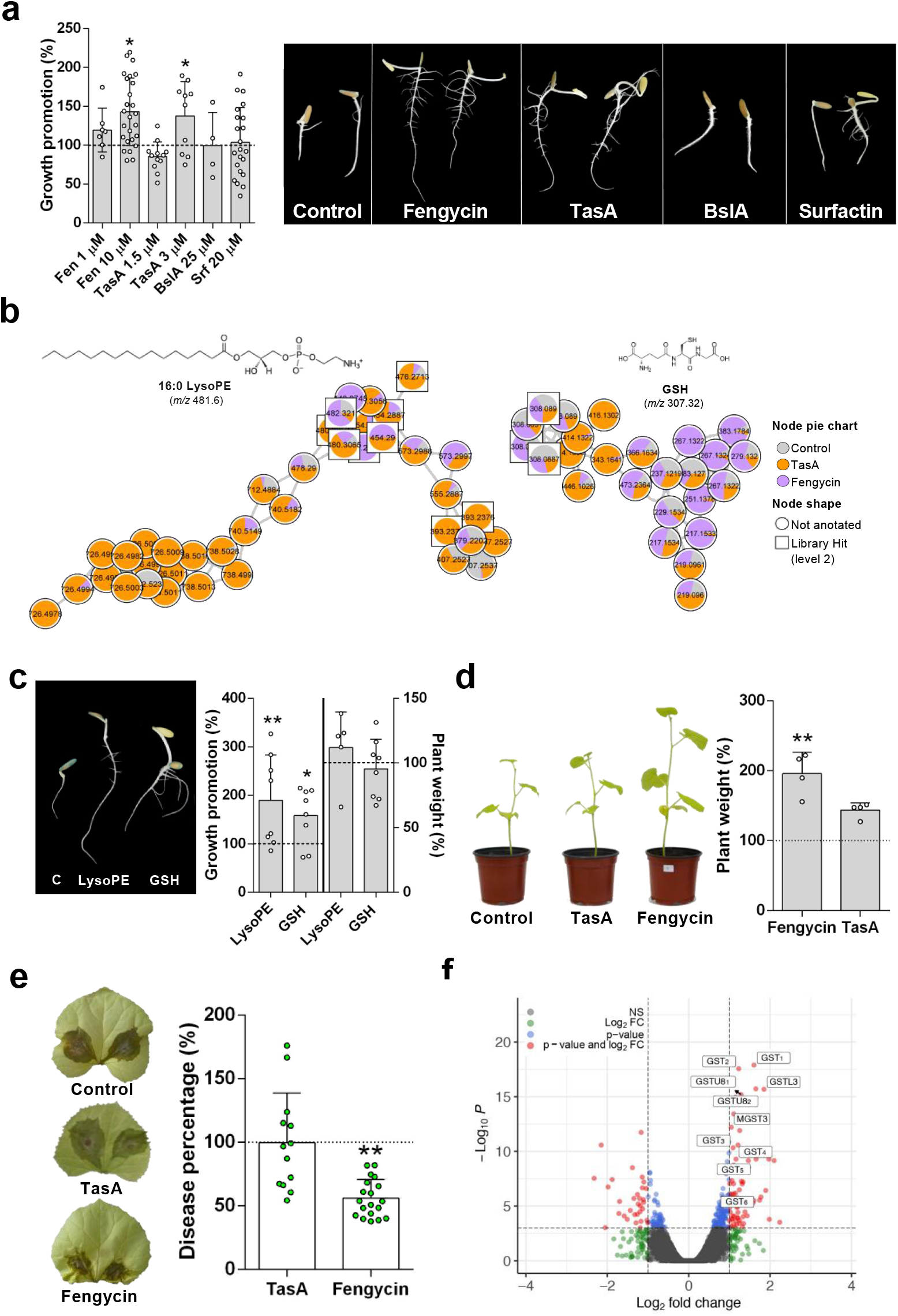
Amyloid TasA and fengycin stimulate radicle development or the growth and immunization of adult plants against aerial necrotrophic fungal pathogens. **A** Left: percentage of an increase in radicle growth of the seed treated with water or purified ECM components normalized to the average radicle area of untreated seeds (100%, discontinued line) five days after the treatments (n= at least 8 except for bslA treatment with n=4). Right: Representative radicles of untreated seeds or seeds treated with purified ECM five days after the treatment. **B** Molecular families corresponding to LysoPE and GSH related to the promoting activity of purified TasA and fengycin based on metabolomic or transcriptomic approaches. The chemical structures of annotated features based on spectral matches to GNPS libraries are also presented for each molecular family. **C** Left: Representative radicles from untreated seeds or seeds treated with LysoPE and GSH five days after the treatment. Right: Left Y axis, percentage of an increase in radicle growth of the seeds treated with water or purified LysoPE and GSH normalized to the average of untreated radicle area (100%, discontinued line) five days after the treatment (n= at least 8); right Y axis, the percentage of weight of adult plants grown from the seeds treated with LysoPE and GSH normalized to the weight of the plants grown from untreated seeds (100%, discontinued line) (n= at least 5). **D** Left: Representative adult plants grown from seeds treated with water (control), 10 μM fengycin, or 3 μM TasA. Right: The percentage of the height of the plants grown from the seeds treated with vehicle control, 10 μM fengycin, or 3 μM TasA normalized to the average height of the plants grown from the control seeds (100%, discontinued line) (n=4). **E** Left: Development of necrotic symptoms in the leaves of adult plants grown from the seeds treated with water (Control), 10 μM fengycin, or 3 μM TasA 72 hours after the treatment with *Botrytis cinerea* spores. Right: Percentage of disease symptoms calculated by measuring the lesion areas normalized to the average lesion area of the control samples (100%, discontinued line) (n= at least 13). **F** Volcano plot representation of DEGs identified by total transcriptome analysis in the leaves of the plants grown from control seeds or seeds treated with fengycin 48 hours after the treatment. Tags label the genes related to glutathione metabolism: GST_1_ (glutathione S-transferase), GST_2_ (glutathione S-transferase), GSTL3 (glutathione S-transferase L3-like), GSTU8_1_ (glutathione S-transferase U8-like), GSTU8_2_ (glutathione S-transferase U8-like), MGST3 (microsomal glutathione S-transferase 3), GST_3_ (glutathione S-transferase), GST_4_ (glutathione S-transferase), GST_5_ (glutathione S-transferase), and GST_6_ (glutathione S-transferase). Average values are shown, and error bars represent SD. Statistical significance was assessed by one-way ANOVA with *post hoc* Dunnett’s multiple comparisons test (each treatment vs. control treatment) in panels **A, C, D**, and **E**.

The total metabolome of radicles emerged from the seeds treated with fengycin or TasA was analyzed to define the contribution to the metabolic signatures associated with the promotion of radicle growth. The results of PCA and heatmap analyses clearly indicated sample clustering in three groups based on metabolomic signatures (Extended Data Fig. 8a, b). Statistical analysis by partial least-squares discriminant analysis (PLS-DA) indicated that glycerophospholipids, fatty acyls, organooxygen compounds, or carboxylic acids were discriminating metabolites in the case of TasA or fengycin treatments, and FBMN of these features showed integration into molecular families composed of other metabolites that generally followed the same abundance patterns (Extended Data Fig. 8c, d, e). Further refinement of this analysis identified two molecular families of interest: i) containing lysophosphatidylethanolamines (LysoPE) and mainly associated with TasA treatment and ii) containing an analog of reduced glutathione (GSH) that accumulated in the radicles grown from the seeds treated with fengycin (Fig. 3B).

Glycerophospholipids play both structural and signaling roles in the plants, and this bifunctionality is in part due to continuous synthesis and turnover of endogenous pools^40^. Accumulating evidence suggests an expanding role of lysophospholipid derivatives in signaling processes in the plant cells and in a variety of response mechanisms. LysoPE treatment has been reported to delay fruit softening when used postharvest, mitigate defoliation effects of ethephon, and delay leaf and fruit senescence in tomato and potato^41,42^ and is used commercially as a plant bioregulator to improve plant product quality^40,43^. On the other hand, GSH is an essential metabolite that has antioxidant functions, which is the principal biological property of GSH; GSH is involved in cellular redox homeostasis and plays essential roles in the development, growth, and environmental response of the plants^44^. Thus, the analysis of metabolomic features of the radicles suggested certain roles for glutathione and *lyso*phospholipids in all signaling events triggered in the seeds after the treatment with *Bacillus* and mediated by the action of fengycin and TasA, respectively. To confirm hypothetical involvement of both molecules in the growth-promoting activity, the seeds were treated with commercially available LysoPE or GSH (Fig. 3C). Both treatments enhanced radicle growth similar to the effect observed after seed treatments with fengycin or TasA.

Enhanced area of the radicles developed from the seeds treated with *B. subtilis* cells and a resulting increase in efficiency of absorption of the nutrients is the easiest interpretation explaining the overgrowth of adult plants from the treated seeds. If this hypothesis is true, we anticipated to detect a long-term promoting effect on adult plants developed from the seeds treated with TasA or fengycin. Only plants emerged from the seeds treated with fengycin were notoriously larger and more vigorous than plants grown from untreated or TasA-treated seeds from the initial stages to the end of the experiment (Fig. 3D). Adult plants grown from the seeds treated with LPE or GSH did not show a significant increase in the growth over time compared to the growth of plants from untreated seeds (Fig. 3C). These findings suggested differences between enhanced growth of adult plants and major growth of the seed radicles observed after the promoting treatments, indicating the presence of different signaling events that triggered i) short term radicle growth mediated by TasA, fengycin, or associated LysoPE and GSH molecules and ii) long term overgrowth of adult plants specifically associated with fengycin. GSH associated to treatments to the seed with fengycin did not produce sustained long-term growth-promoting effect, suggesting that constant endogenous trigger of the events is required and can only be achieved by fengycin treatment.

Previous studies have shown that application of fengycin elicits plant defense responses in adult plants^12,45^. Therefore, we investigated whether, in addition to the long-term growth-promoting effect, treatment of the seeds with fengycin immunizes adult plants against aboveground pathogens. The third leaves of adult plants were inoculated with a spore suspension of the necrotrophic fungal pathogen *Botrytis cinerea*. The size of necrotic lesions induced by the fungus in the plants grown from fengycin-treated seeds was significantly smaller than that of the control plants and plants grown from TasA-treated seeds (Fig. 3E). The results of transcriptomic analysis of the leaves indicated that the treatment of the seeds with fengycin induced higher expression of the genes related to glutathione metabolism, specifically including glutathione S-transferases (GSTs), 48 hours after the challenge with *B. cinerea* (Fig. 1F, Extended Data Fig. 9a). Participation of GSTs in antioxidant reactions together with the important cellular antioxidant GSH was clearly shown to mitigate oxidative stress caused by this necrotrophic fungus in infected tissues^46^. Interestingly, metabolomic data obtained from the radicles from fengycin-treated seeds demonstrated an increase in the levels of certain molecules of the same cluster that included GSH (Fig. 3B), and adult plants grown from bacterized seeds accumulated certain molecules belonging to the GSH cluster, especially in the aerial region (Extended Data Fig. 9b). Overall, these results suggested that initial seed treatment with fengycin triggered the accumulation of GSH in the seedlings, which conferred enhanced antioxidant capacity able to mitigate an imbalance in the redox status in adult plants imposed by infection with *B. cinerea*.

### Different promoting activities of TasA and fengycin rely on specific interactions with seed oil bodies

The perisperm-endosperm of the melon seeds is enriched in oil bodies (OBs), which are nutrient reservoir organelles distributed in various plant organs and mainly composed of triacylglycerides (TAGs) and other neutral lipids. During germination, the monolayers of these covered vesicles are degraded during interaction with glyoxysomes, and released contents become available to catabolic routes that feed the embryo^47,48^. Accumulation of bacterial cells in the micropylar endosperm and major changes in the lipid composition of the radicles and adult plants after the treatments suggested that the oil bodies are the target of the growth-promoting molecules fengycin and TasA. The results of confocal laser scanning microscopy (CLSM) of thin sections of the seeds using a neutral lipid stain 16 hours after seed treatment indicated abundant presence of the oil bodies in the endosperm of untreated seeds. In agreement with our hypothesis, we detected certain regions of the endosperm with high levels of disaggregated oil bodies that were always surrounded by bacterial aggregates (Extended Data Fig. 10a).

Treatments with purified TasA or fengycin promoted disaggregation of OBs similar to the effect of the treatments with *Bacillus* cells (Fig. 4A, top). The results of transmission electron microscopy analysis (TEM) of thin sections of negatively stained seeds did not detect significant anatomical differences between the treatments in addition to OB disaggregation except for the localization of OB-surrounding glyoxysomes in samples treated with TasA, indicating the presence of interacting organelles where oil bodies are degraded to release stored energy^48,49^ (Extended Data Fig. 10b, c). OBs purified from melon leaves were treated with TasA or fengycin and double stained with Fast Green FCF to stain the proteins and with Nile red (Fig. 4A, bottom). The results of CLSM analysis indicated certain differences in the size and pattern of aggregation of OBs in various treatment groups; fengycin induced disaggregation of OBs observed *in vivo*, and TasA preferentially induced a higher level of aggregation of individual OBs than that observed *in vivo* (Extended Data Fig. 10d). Higher intensity of the green fluorescence signal near OB aggregates in the samples treated with TasA suggested nonspecific staining of the amyloid protein and its localization around the vesicles. The results of TEM analysis of purified OBs and immunogold labeling with anti-TasA antibodies indicated the presence of gold particles decorating the perimeter of the OBs and interconnecting TasA fibers (Extended Data Fig. 10e). Thus, we hypothesized that the effect of TasA involves interactions with structural proteins present on the surface of the OBs, which enhances association of OBs with glyoxysomes and/or other degradative processes that eventually lead to specific release of the lipids. For example, accumulation of *lyso*phospholipids correlated with growth promotion of seed radicles but did not correlate with long-term effects in adult plants. The monolayer of OBs mainly contains oleosins, which are structural proteins that modulate the stability and size of OBs, regulate lipid metabolism, and play an important role in OB degradation^50–52^. The results of the pulldown assays of whole protein extracts of the seeds indicated coelution of purified TasA with oleosin (Fig. 4B and Table S5). Interaction of the two proteins was confirmed by far-western blot analysis of purified TasA and purified oleosin 1 from *Arabidopsis thaliana* (Fig. 4C). These results suggested that the TasA-oleosin interaction was responsible for the aggregation of OBs *in vitro* and most likely accounted for the accumulation of *lyso*phospholipids, which are the signaling molecules related to short-term growth promotion of the radicles. Modulation of activity of oleosin as a regulator of lipid catabolism or the level of the degradation of the lipids are alternative but not exclusive explanations for this process mediated by TasA.

**Figure 4.**
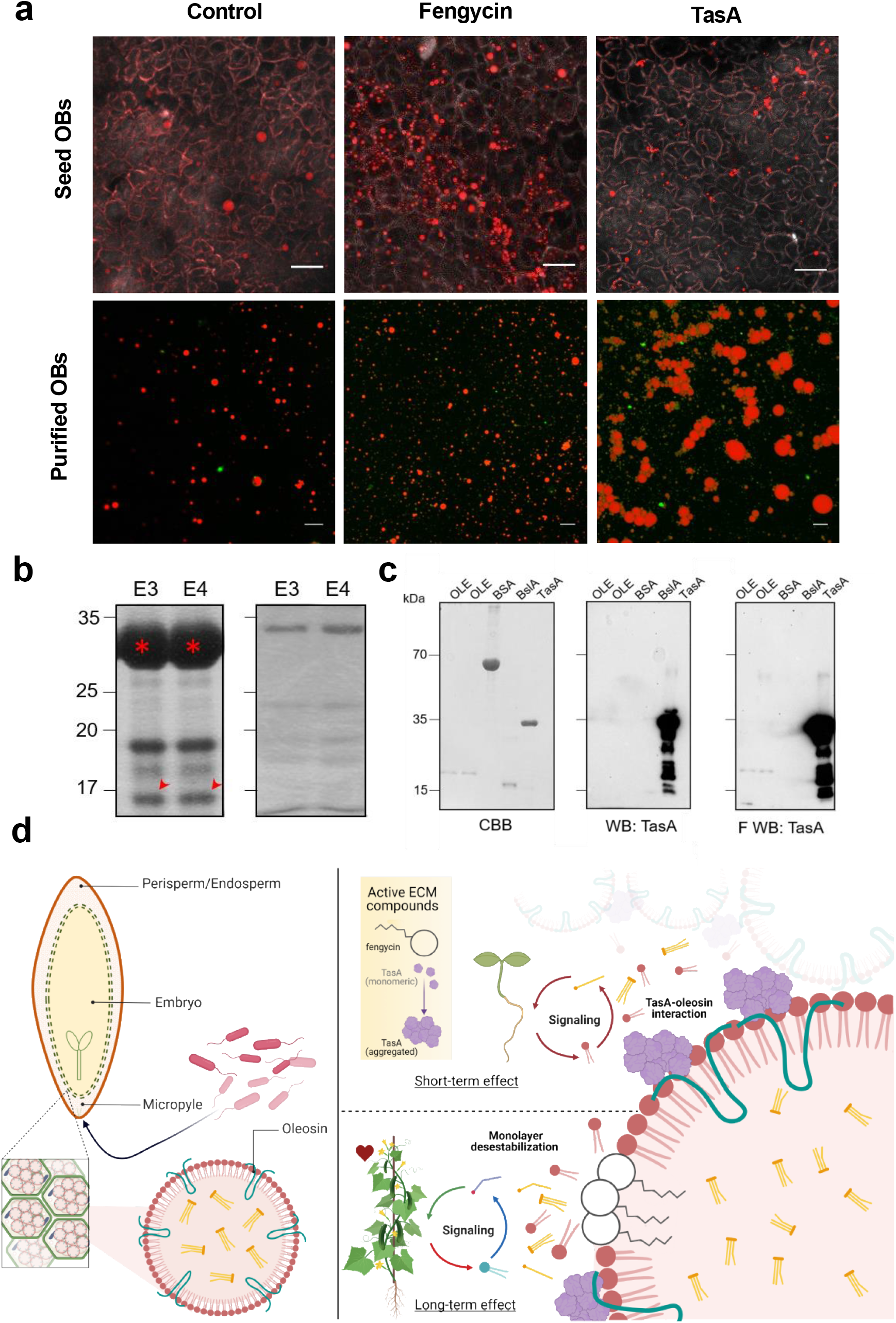
Differential interactions of amyloid TasA and fengycin with oil bodies are related to specific stimulation of plant growth. **A** CLSM images of transversally cut seeds 16 hours after the treatment with 10 μM fengycin or 3 μM TasA (top; scale bar: 20 μm) and purified oil body suspension 16 hours after the addition of 10 μM fengycin or 3 μM TasA (bottom; scale bar: 5 μm). Oil bodies were stained with Nile red, and membrane proteins were stained with Fast Green FCF. **B** Coomassie brilliant blue (CBB)-stained SDS-PAGE gel of elution fractions of pulldown assay of TasA with proteins in the extracts from melon seeds. The coeluting band is marked with red arrows and identified as an oleosin by HPLC-ESI-MS/MS analysis. **C** Far-Western blotting: left, a CBB-stained SDS-PAGE gel of fractions of oleosin obtained during purification; BSA was used as a control, and fractions of BslA and TasA were obtained during purification; middle, immunoblot of purified proteins using an anti-TasA antibody (1:20,000); right, far-western blot using an anti-TasA antibody (1:20,000) after renaturing of the proteins and incubation with TasA protein as a bait (right). **D** Overall scheme of the proposed mechanism of growth-promoting effects of TasA and fengycin. Left: Structure of melon seeds and oil bodies (OB). The perisperm/endosperm of the seeds is enriched in storage cells containing abundant OBs. OBs are composed of a monolayer of phospholipids and structural proteins, mainly oleosins, and contain neutral lipids, such as triacylglycerides. After the treatment, *B. subtilis* cells passively enter the inner tissues of the seeds. Right: Inside the seeds, aggregates of TasA interact with oleosins of OBs, triggering the accumulation of lysophospholipids involved in signaling related to a short-term effect (radicle growth). Fengycin destabilizes the OB membrane, and the perturbation triggers the accumulation of specific molecules, such as GSH, increasing the initial pool of this antioxidant molecule implicated in long-term stimulation of adult plant growth and immunization.

Fengycin has been described to efficiently interact and disrupt artificial lipid monolayers or bilayer membranes in a concentration-dependent manner, which also modulates the level of aggregation of the lipopeptide^53–55^. Depending on the fengycin concentration, two mechanisms of action have been proposed: i) low concentrations promote the formation of the aggregates that induce the formation of the pores and subsequent changes in membrane permeability; and ii) high concentrations cause solubilization of the membranes similar to the effect of detergents^53,54^. In addition to micellar concentrations of fengycin, specific disruption of the membranes relies on the lipid composition of the target^56–58^. Exact chemical composition of phospholipids of the membranes of OBs in the seeds is not known and varies between the species^59^; however, we propose that high affinity of fengycin for lipid membranes explains disaggregation and a reduction in the size of OBs. As a consequence of alterations in the OB membranes, endosperm cells may overproduce GSH, leading to an increase in the basal level of this molecule detected by metabolomic analyses in the present study, and may account for long-term effects of fengycin treatment of the seeds.

These findings indicated that beneficial effect of seed treatment with *B. subtilis* on the melon plants was associated with changes in the contents and specific pools of metabolites released from the storage tissues, which were mediated by at least two molecules, fengycin and TasA. We propose that specific interaction of TasA or fengycin with OBs defines two different physiological responses: i) the promotion of radicle growth mediated at least by the accumulation of lysophospholipids and ii) overgrowth of adult plants and immunization against aerial pathogens mediated by the accumulation of various molecules, such as GSH, specifically elicited by fengycin (Fig. 4D). Thus, the proposed mechanism of action of these two molecules depends on the internal structure of the seeds, which directly influences the accessibility of the molecules in the storage tissues, which contain abundant OBs, and initial lipid composition of the seeds, which modulates specific release of the signaling molecules. We propose that only seeds containing abundant OBs and characterized by specific morphology of the primordial tissues respond to the beneficial interaction with *Bacillus* mediated at least by fengycin and TasA. According to this model, wheat or maize, which are monocotyledonous plants whose seeds are composed of starchy endosperm surrounded by a layer of living cells, did not react to the presence of fengycin, and the treatment of *Cucumis sativus* or soybean seeds, which are anatomically different but contain OBs, induced a growth-promoting phenotype (Table S6).

## Acknowledgements

We thank Saray Morales Rojas for technical support, Josefa Gómez Maldonado from the Ultrasequencing Unit of the SCBI-UMA for RNA sequencing, Casimiro Cárdenas for protein analysis (Proteomic Unit, SCAI-UMA), Juan Félix López Téllez for technical support in the transmission electron microscopy analysis and John Pearson for his technical support in the confocal laser scanning microscopy analysis (Bionand). This work was supported by grants from ERC Starting Grant (BacBio 637971) Plan Nacional de I+D+i of Ministerio de Economía y Competitividad and Ministerio de Ciencia e Innovación (AGL2016-78662-R and PID2019-107724GB-I00) and Proyectos I+D+I en el marco del Programa Operativo FEDER Andalucía (UMA18-FEDERJA-055). M.V.B. is the recipient of an FPU contract (FPU17/03874) from the Ministerio de Ciencia, Innovación y Universidades. C.M.S is funded by the program Juan de la Cierva Incorporación (IJC2018-036923-I). D.P. was supported by the German Research Foundation (DFG) with Grant PE 2600/1. AMCR and PCD were supported by the National Institutes of Health (NIH) grant DP2GM137413. PCD was supported by the Gordon and Betty Moore Foundation through Grant GBMF7622, the U.S. National Institutes of Health for the Center (P41 GM103484, R03 CA211211, R01 GM107550).

## Author Contributions

**DR**, conceived, designed the work, drafted and edited the text; **MVB**, collected most of experimental data, and drafted the manuscript; **CMS,** designed, collected and analyzed MS data, and edited the manuscript; **AMC**; collected and analyzed MS data, and edited the text; **DP**, collected MS data, and edited the text; **LD**; informatically analyzed data and drafted figures; **AdV**, substantially revised and edited the text; **AP**; substantially revised and edited the text; **PCD**, substantially revised and edited the text.

## Notes

### Competing Interest Statement

The authors have declared no competing interest.

